# Enemy mitigation in farmlands: Intensive agriculture impacts avian nesting and subsequent ectoparasite infestation

**DOI:** 10.64898/2026.02.02.703377

**Authors:** Simon Coroller-Chouraki, Allen Bush-Beaupré, Jade Savage, Marc Bélisle

## Abstract

Intensive agricultural practices directly affect farmland bird and non-target insect populations by modifying their habitats, but may also act indirectly by altering their interactions. Notably, the breeding success of insectivorous birds has been shown to suffer from reduced prey availability. Yet little is known about how agriculture influences host-parasite relationships in wild birds. How agricultural intensity affects parasites, and whether this alleviates or exacerbates the trophic stress imposed on birds therefore remains to be determined. We estimated the number of obligate hematophagous *Protocalliphora* blowfly larva (Diptera: Calliphoridae) that parasitized nestlings in 2,560 Tree Swallow (*Tachycineta bicolor*) broods along a 10,200-km² gradient of agricultural intensity between 2004 and 2019 in Québec, Canada. We first modeled two key variables along the causal paths expected to affect *Protocalliphora* prevalence and load (abundance) within infested broods: nestling hatching date and nestling host availability. Hatching phenology varied by several days with early-spring meteorological conditions and parental age, as for nestling availability (nestling-days), which also decreased along the agriculture intensity gradient as pastures and hay fields were replaced by large-scale, cereal row crops. Nestling availability peaked under low precipitation rates when temperatures reached 18 to 25 °C. Prevalence and load of blowfly larvae directly increased with nestling availability as well as with the temperature and precipitation that occurred during the larval development and pupation stages. Controlling for nestling availability, *Protocalliphora* prevalence and load peaked in forested landscapes interspersed by pastures and hay fields and reached their lowest in landscapes dominated by corn and soybean monocultures with minimal tree cover. Agricultural intensity thus reduced infestation likelihood and severity both directly and indirectly, by limiting nestling host availability. This finding is notable given the documented negative effects of agricultural intensity on fledgling number and body condition in farmland birds, even after controlling for insect prey reduction. If agricultural intensity indeed reduces the parasitic pressure exerted by bird blowflies and its consequences for fledgling condition and recruitment, this suggests that other agricultural impacts (e.g., toxicological effects from pesticides) may play a larger role than previously recognized in the severe declines of farmland bird populations observed across the Holarctic.

**Open research statement:** The data supporting this study are not yet publicly available, as they require final harmonization, documentation and anonymization prior to archiving. Upon acceptance of the manuscript, all underlying data and associated code will be permanently deposited in the Zenodo repository and made fully accessible with a DOI.

## Introduction

Agricultural intensification has been acknowledged as a major driver of the changes in structure and dynamics that farmland bird and insect populations have experienced over the last few decades (Stanton et al. 2018, Spiller and Dettmers 2019, Raven and Wagner 2021, Outhwaite et al. 2022, Rigal et al. 2023). For instance, the breeding season emerges as a pivotal period for farmland bird populations, as offspring production and survival combined with immigration can buffer losses due to emigration or mortality, albeit at the potential expense of substantial parental care (Hanssen et al. 2005, Newton 2013, Le Vaillant et al. 2026). Yet, during this critical period, adverse agricultural practices, such as large-scale row-cropping with simplified crop rotations (Benton et al. 2003), pesticide use (Mineau and Whiteside 2013, Gibbons et al. 2015, Moreau et al. 2022, Molenaar et al. 2024), and faster harvesting and mowing regimes (Tews et al. 2013, Bretagnolle et al. 2018) directly affect bird abundance and fitness. These components of modern agricultural practices also affect insect communities (Forister et al. 2019, Uhl and Brühl 2019, Raven and Wagner 2021, Pawan Kumar et al. 2023) and thereby trophic interactions among insects and birds (Emmerson et al. 2016, Rigal et al. 2023).

Most farmland birds are insectivorous during the breeding season and exert top-down pressures on insects (Mäntylä et al. 2011, Bucher et al. 2019). Reductions in insect availability and/or nutritive quality due to agricultural intensification can thus lead to lower reproductive success in birds (Twining et al. 2016, Garrett et al. 2022c, Grames et al. 2023, Goebel et al. 2024). Analogously, some arthropods found in agroecosystems, such as bird lice (Phthiraptera), *Dermanyssus* mites (Mesostigmata: Dermanyssidae), some *Ixodes* ticks (Ixodida: Ixodidae), and *Protocalliphora* blowflies (Diptera: Calliphoridae), are obligate ectoparasites that rely on avian nesting and host density to thrive at their expense (Arneberg et al. 1998). While a few studies addressed the potential cumulative impacts of agricultural intensity and ectoparasite load on avian fitness in agroecosystems (e.g., Sigouin et al. 2021), how agricultural intensity may directly and indirectly affect ectoparasite loads of farmland birds remains largely unexplored (Dietsch 2005, Daoust et al. 2012a). Indeed, besides studies on small mammals (Shilereyo et al. 2022), lizards (Biaggini et al. 2009), and poultry (e.g., Salam et al. 2009, Sparagano et al. 2014, Mata et al. 2018, Chambless et al. 2022), we found only one study that formally addressed variation in wild bird ectoparasite loads across a gradient of agricultural intensity (Daoust et al., 2012a). This study, which spanned two years of sampling, showed that nests of Tree Swallows (*Tachycineta bicolor)* bore lower loads of *Protocalliphora* blowfly larvae in farmlands dominated by intensive, cereal row-cropping than in farmlands focusing on forage production. This lack of research on the effects of anthropogenic pressures on *Protocalliphora* loads (Eeva et al. 1994, Gentes et al. 2007, Eeva and Klemola 2013) stands in stark contrasts to the substantial focus on their potential impacts on their hosts’ fitness (Maziarz et al. 2022).

Aside of the direct effects that habitat loss and pesticides may have on ectoparasites (Bennett and Whitworth 1991, Daoust et al. 2012a), agrointensive practices may affect the ectoparasite loads of farmland birds through diverse pathways. For instance, intensive agriculture may reduce avian host density and diversity at the landscape scale and thereby limit *Protocalliphora* blowfly population sizes (Bennett and Whitworth 1992, Arneberg et al. 1998, Stanton et al. 2018). Agricultural intensity may further reduce “local” host abundance and temporal availability for *Protocalliphora*, by decreasing the clutch size, hatching rate, and nestling survival of farmland birds (Stanton et al. 2018, Bretagnolle et al. 2018, Garrett et al. 2022b, Grames et al. 2023, Rigal et al. 2023, Goebel et al. 2024). Lastly, agricultural intensity may affect *Protocalliphora* loads by altering the quality of their farmland bird hosts (e.g., via their contamination by pesticides and immunological response, Sigouin et al., 2021) or the community of their natural enemies (i.e., predators and hyperparasites, (Daoust et al. 2012a, Schöfer et al. 2023, Sulg et al. 2023).

Measuring the influence of agricultural intensity on ectoparasite loads of farmland birds is thus crucial to properly interpret the effect, or lack thereof, of agriculture on avian fitness (Thomas et al. 2007). Indeed, the impacts of some intensive agricultural practices on farmland birds may manifest and even be amplified through increased ectoparasite loads or be alleviated if such practices reduce ectoparasite loads. The debated negative impacts of *Protocalliphora* blowflies on avian nestling development and survival induced by blood loss and infection risks (Williams 2017) likely exemplifies the outcome of such intricated effects of environmental conditions on host-parasite dynamics. For instance, conflicting results have been reported regarding nestling tissue maturation and physiological parameters (Johnson and Albrecht 1993). While some studies reported fatal threats to nestlings (Merino and Potti 1996, Bańbura et al. 2004), others found neutral (Thomas et al. 2007) or even positive associations of *Protocalliphora* loads with nestling growth rate and fledging success (Musgrave et al., 2019), possibly induced by parental care. These studies, however, involved different bird and *Protocalliphora* species, with potential varying susceptibility levels and impacts, and were conducted across diverse regions with distinct environmental conditions. Moreover, the full extent of *Protocalliphora* blowflies’ impact may not be apparent within the nest and may manifest itself only later through diminished post-fledging behavior and survival (Thomas et al. 2007, Streby et al. 2009). More importantly, these blood-feeding ectoparasites are hypothesized to exacerbate the effects of environmental stressors such as food scarcity, inclement weather, and contaminants (Sabrosky et al. 1989, Howe 1992, Merino and Potti 1996, Musgrave et al., 2019, Sigouin et al. 2021), many of which may be influenced by agricultural intensity and show interacting effects with the latter (Garrett et al. 2022a,b).

As a first step, the use of a causal inference framework applied to longitudinal data collected from many sites distributed over a large spatial scale could provide valuable insights into the complex relationships linking agricultural intensification and other environmental contexts to farmland bird nesting events and ultimately, their *Protocalliphora* parasitic load. Such analysis requires identifying a plausible causal structure linking relevant variables and controlling for the proper sets of variables to avoid biases when assessing specific effects of agricultural intensification (Cinelli et al. 2022, Arif and MacNeil 2023). Prior research found that several meteorological factors influence *Protocalliphora* loads in bird nests. For example, higher loads have been associated with temperatures ranging between 23 and 25 °C, and lower loads with higher humidity levels (Dawson et al. 2005, Mennerat et al. 2021, Maziarz et al. 2022). Nonetheless, weather-load associations vary significantly across studies (e.g., Musgrave et al. 2019, García-del Río et al. 2024), notably as temperature and precipitation can have interacting effects and because studies have been conducted under contrasting climates (Merino et al. 2024). Furthermore, potential effects of early-spring meteorological contexts on *Protocalliphora* have so far been neglected (but see Mennerat et al. 2021) even though they are known to affect the breeding phenology of birds and thereby *Protocalliphora* host species availability in terms of broods and nestlings within broods (de Zwaan et al. 2022, Imlay et al. 2019, Taff and Shipley 2023). This could explain why *Protocalliphora* loads have been found to both increase (Eeva et al. 1994, Tomás et al. 2007, Wolfe-Merritt et al. 2022) and decrease as the bird breeding season progressed (Wesołowski 2001).

Our understanding of host-parasite associations is often limited by a lack of knowledge about the influence of geography and landscape structure on host and parasite densities, as well as on the dynamics of their interactions (Wells et al. 2015). Here we investigate the intricate causal pathways of direct and indirect effects by which agricultural intensity and meteorological conditions may affect the nesting events of Tree Swallows and their *Protocalliphora* parasitic loads. More specifically, we address how landscape habitat composition and weather (i.e., temperature and precipitation) may affect the prevalence and intensity of Tree Swallow brood infestation by *Protocalliphora,* and this, through their influence on the hatching date and brood size of this secondary cavity-nesting bird (Fig.1). Our measures of direct and indirect effects are based on 2,560 Tree Swallows nesting events, whose nests were examined for *Protocalliphora* pupal remains, that occurred between 2004 and 2019 across a 10,200-km² gradient of agricultural intensity in southern Québec, Canada. By investigating the overlooked interactions between birds and their ectoparasites within agroecosystems, our research advances understanding of how agricultural practices influence farmland birds and their associated communities.

**Figure 1.**
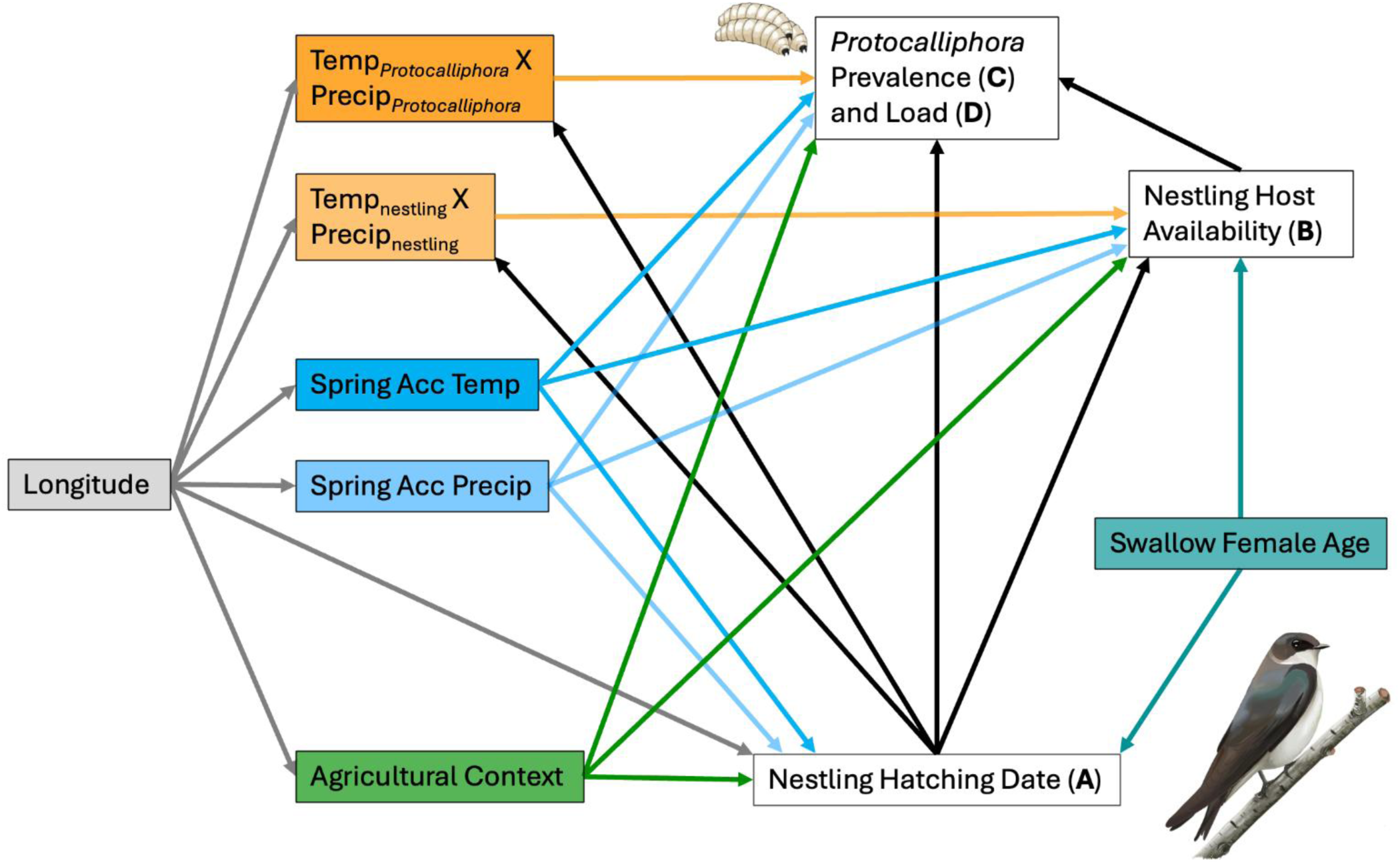
Directed Acyclic Graph of putative causal relationships among studied variables. Direct effects of variables on the first hatching date (A) and nestling host availability (B) provided by Tree Swallow broods were assessed sequentially before estimating those on the prevalence (C) and load (D) of *Protocalliphora* blowfly infestations (see models in Table 1). Spring weather conditions were defined by accumulated daily temperatures (Temp) and precipitation (Prec) between 1 April and 1 June for both swallows and blowflies. Weather conditions during the nestling and larval/pupation phases corresponded to hourly averages over the respective period during which swallow nestlings and blowfly larvae and pupae developed in a given swallow brood (i.e., up to fledging or death and to estimated emergence, respectively). Agricultural context surrounding each swallow brood was defined based on the scores of the first two axes of a robust, compositional PCA fitted on relative land cover values measured annually within 500 m of each nest box of the study system (i.e., Comp.1 and Comp.2 in Fig. 2B).

**Table 1.**
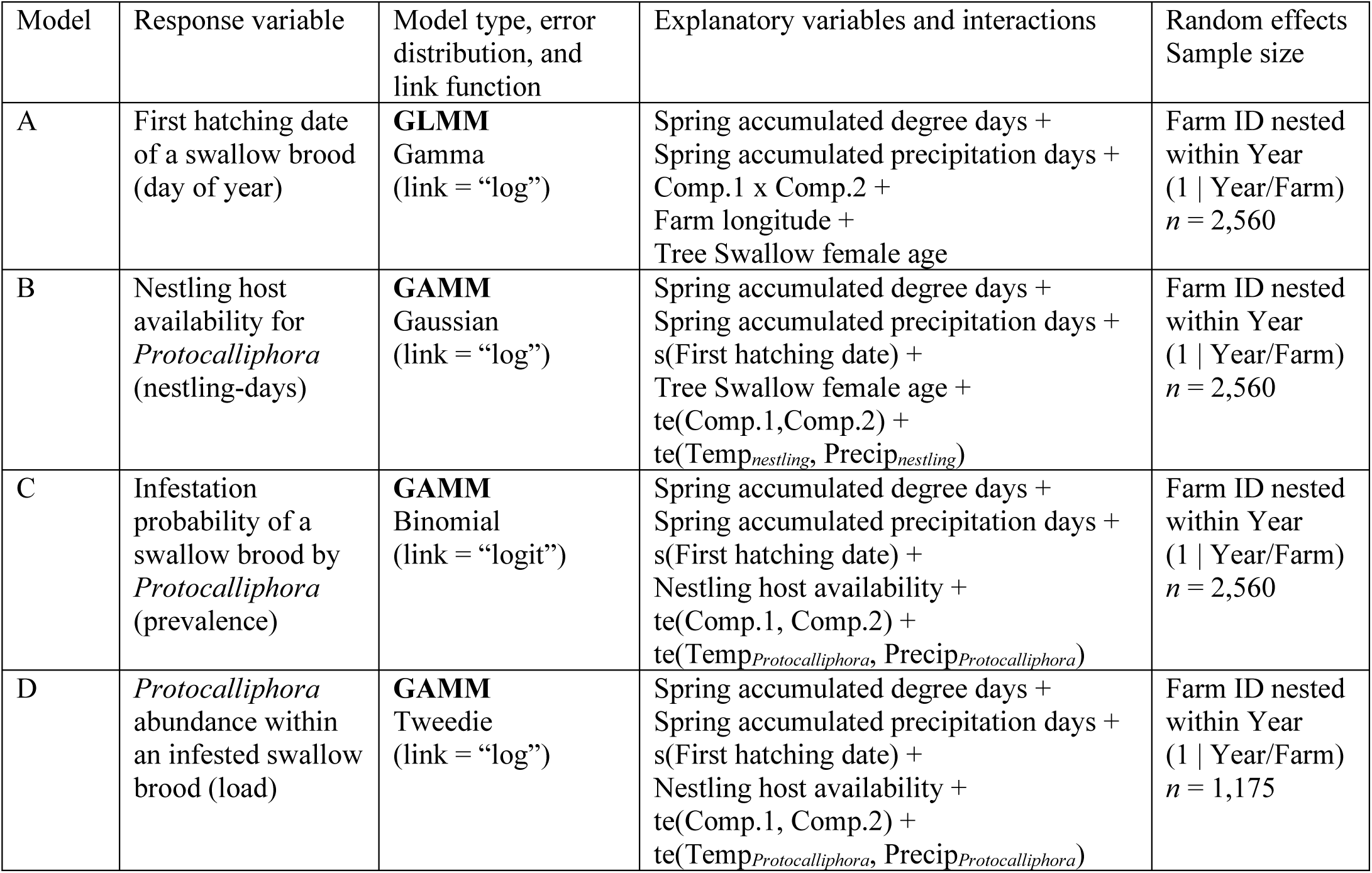
Models used to estimate the direct effects of ecological determinants of *Protocalliphora* blowfly prevalence and load within infested Tree Swallow broods. Models are based on the putative causal structure given by the Directed Acyclic Graph in Fig. 1. For GAMMs, s() and te() refer to simple and full tensor product smooth terms in mgcv, respectively (Wood 2017). Model outputs are provided in Section S1.

## Methods

### Study system and landscape characterization

Our study system was established in 2004 to investigate the effects of agricultural intensification on the breeding ecology of Tree Swallows, an aerial insectivorous passerine bird that readily breed in nest boxes (Ghilain and Bélisle 2008). The system comprised 400 nest boxes distributed equally among 40 farms, located along an east-west gradient of agricultural intensity covering 10,200 km² in southern Québec, Canada (Fig. 2A). Whereas the agroecosystems of the east are dominated by pastures and forage crops embedded into substantial forest, those in the west consist of large expanses of annual row crop monocultures mostly focused on maize, soybean, and wheat associated with extensive pesticide use (Giroux, 2022). Although a few bird species benefitted from the intensification of agricultural practices in the western part of the study area, most farmland bird species have shown decreasing breeding populations in the past three decades (Robert et al. 2019). Nest boxes were typically set 50 m apart along a linear transect that followed a field edge. The entrance of boxes faced the south-east about 1.5 m above the ground.

**Figure 2.**
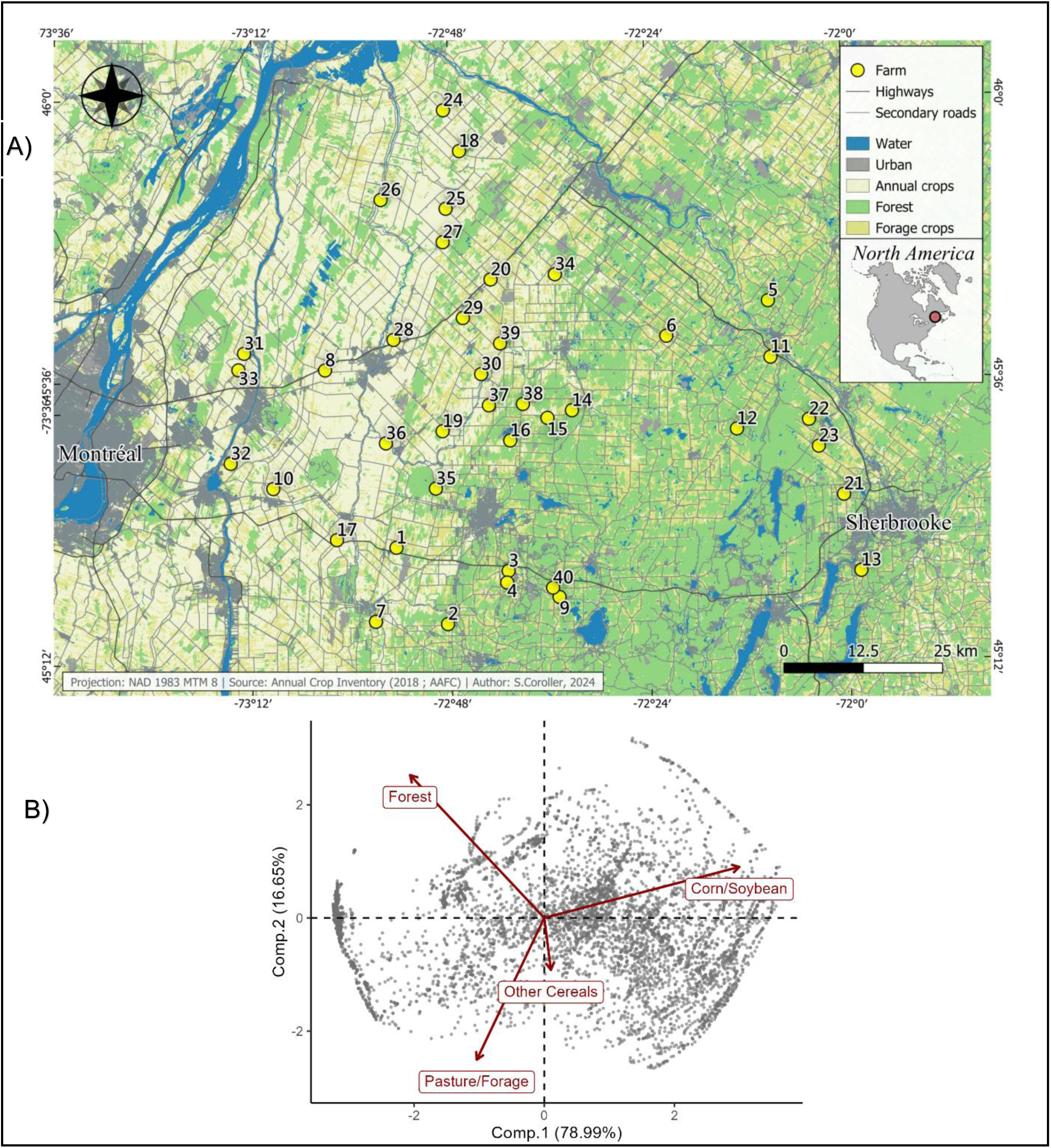
A) Tree Swallow breeding ecology and *Protocalliphora* blowfly infestations were monitored between 2004 and 2019 within a nest-box network composed of 40 farms (10 nest boxes per farm) distributed across a 10,200-km² gradient of agricultural intensification in southern Québec, Canada. Map coordinates follow the WGS 84, EPSG:4326 reference system. Spatial data were projected in the NAD83 MTM Zone 8 system for mapping and spatial analyses. B) Biplot of the robust, compositional PCA fitted on the relative land cover values determined annually around each of the 400 nest boxes (500-m radius) of the study system between 2006 and 2019 (*n* = 5,600 agricultural contexts).

We assessed landscape composition within a 500-m radius of each nest box in our study system by conducting an in-situ characterization of habitat types at the end of each breeding season between 2006 and 2019. Habitat types included “Forest”, “Corn/Soybean”, “Other Cereals” (i.e., wheat, oat, barley), and "Pasture/Forage" (i.e., hay, alfalfa, clover). We used orthophotos (1:40,000) in QGIS (QGIS, 2020) to delineate habitat and agricultural fields boundaries. Following Garrett et al. (2022a) and Powell et al. (2024), we then performed a robust compositional Principal Component Analysis (rPCA) using the robCompositions R package (Templ et al. 2011) on the yearly relative areas covered by the above habitat types around each nest box between 2006 and 2019 (Fig. 2B). Note that streams and water bodies covered on average < 1% of nest box surroundings (Garrett et al. 2022b) and were thus not considered in this analysis. The first component (Comp.1) of the rPCA explained 78.99% of the variance in landscape composition and was positively correlated to maize and soybean cover and negatively to forage and tree covers. The second component (Comp.2) explained 16.65% of the variance in landscape composition and was positively correlated to tree cover and negatively to forage. Because in-situ characterization of crop covers could not be conducted in 2004 and 2005, we had to extrapolate Comp.1 and Comp.2 score values for each nest box on each of these two years. We did this by averaging score values from 2006 to 2008 for each nest box, and assigning these values to both 2004 and 2005, given that the habitat composition of a landscape varied only marginally between years (Figure S1). This procedure ensured that each nest box was assigned a yearly landscape context defined by Comp.1 and Comp.2 score values. Landscape contexts based on crop covers like the above were found to be associated to various aspects of the breeding phenology (Bourret et al. 2015, Courtois et al. 2021) and success (Garrett et al. 2022a,b,c) of Tree Swallows in our study area.

### Swallow and *Protocalliphora* basic biology and data

Whereas adult *Protocalliphora* feed on nectar or pollen, their larvae are obligate hematophagous parasites of nestling birds (Bennett and Whitworth 1991). Female adult flies locate avian nests using visual and olfactory cues (Williams 2017), and lay up to 90 eggs on or next to the bird nestlings (Sabrosky et al. 1989). *Protocalliphora* larvae reside at the bottom of the nest, migrating up only to feed on the blood of nestlings. If *Protocalliphora* larvae typically attach to legs, abdomen, or rump, they sometimes target nasal and auditory cavities, potentially leading to infections or deformities (Streby et al. 2009). Over 7 to 15 days, *Protocalliphora* larvae will ingest 3 to 4 blood meals before pupating among nest material towards the nest bottom (Bennett & Whitworth, 1992). Nestlings dying or leaving the nest rapidly may hence act as a potential time limiting factor for *Protocalliphora* larval. Adult flies emerge after nestlings have either perished or fledged, leaving the remains of their puparia within the nest material (Jánošková et al. 2010). *Protocalliphora* are univoltine and overwinter as adults before reproducing the following spring (Gold and Dahlsten 1983, Bennett and Whitworth 1991).

Each year between 2004 and 2019, we collected the content of all nest boxes that had been occupied by Tree Swallows once they had left the study area and initiated their fall migration. Nests were then stored in plastic bags at a temperature of 4 °C until the day before processing, when they were stored at −80 °C to kill any remaining organisms. Nests were processed under a ventilated hood to retrieve all *Protocalliphora* puparia. We could only retrieve puparia as maggots that did not reach the pupal stage had decomposed by then. Our measurements of *Protocalliphora* occurrence and abundance may thus be underestimated because of this survival bias (Jánošková et al. 2010). That said, we expect this bias to be slight as underfed *Protocalliphora* larvae can pupate as "runts" (Sabrosky et al. 1989, Bennett and Whitworth 1991). We restricted this paper’s dataset to nest boxes with complete environmental information (i.e., with landscape, swallow breeding phenology and success, as well as weather data) and excluded all nest boxes that hosted more than a single bird clutch on a given year to ensure that collected *Protocalliphora* puparia did not origin from multiple bird broods. Although three species of *Protocalliphora* have been found to parasite Tree Swallows in variable proportions across swallow broods, farms and years in our study system (Daoust et al. 2012b, Coroller-Chouraki et al. 2026), one species, namely *P. sialia*, clearly dominates with 80.6% of subsampled puparia (*P. bennetti*: 15.8%, *P. metallica*: 3.6%; *n* = 3,009 puparia from 216 Tree Swallow nests). Since *P. sialia* has remained dominant over the years (i.e., 61.7% to 98.4% of subsampled puparia), that the three *Protocalliphora* species are similar in larval size and behavior, and that it is virtually impossible to identify all puparia by species, we have chosen to treat parasitic events by *Protocalliphora* without regard to their species composition.

We monitored nest boxes every two days to document the phenology (nest building, egg laying, incubation, hatching, fledging) and success (clutch and brood sizes, growth and fate of each nestling) of each Tree Swallow breeding event. We captured breeding females during incubation and aged them [as second year (SY) or after second year (ASY)] based on either plumage or band number, if marked as nestlings in our system (see Garrett et al. 2022b for details). Female age is a key determinant of breeding phenology and success in Tree Swallows (Courtois et al. 2021). Because female *Protocalliphora* lay their eggs in nests with born nestlings, we used the ordinal date on which the first egg of a given swallow clutch hatched to assess how phenological differences among swallow broods affect the occurrence and abundance of *Protocalliphora* within swallow nests. The timing of swallow breeding events will notably affect the number of host nests available to *Protocalliphora*, which can prompt nest infestation by the flies while simultaneously diluting infestation risk and severity among nests. It is also associated to weather and within-nest availability of nestlings and may thereby affect the extent to which *Protocalliphora* parasite a given swallow brood and the conditions under which their eggs and larvae will develop. The time window during which nestlings of a given swallow brood were exposed to weather conditions (see next section) and available to *Protocalliphora* parasitism thus started when its first egg hatched and ended when its last nestling either fledged or died. We used the number of nestling-days during this period (by summing the number of days each nestling was alive in the nest box) to estimate nestling availability to *Protocalliphora* for each brood. We extended the nestling availability period by seven days (Sabrosky et al. 1989) when estimating the weather conditions experienced by *Protocalliphora,* from eggs to emergence, in each individual nest box (see next section).

### Meteorological data

Spring meteorological conditions, and notably temperature and precipitation summarized in terms of accumulated degree days (ADD) or accumulated precipitation days (APD), respectively, have been found to affect the timing of egg-laying and clutch size in birds (Bourret et al. 2015, Sockman and Courter 2018, Simmonds 2019), as well as the phenology and magnitude of prey availability for breeding insectivorous birds (Bolduc et al. 2013, Belitz et al. 2025). Following agronomical prescriptions for our study area (Agrométéo, 2024), we computed ADD for each farm on each year by first averaging the minimum and maximum temperatures (°C) of each day between 1 April (day of year, DOY, 91) and 1 June (DOY 152; start of the hatching period; “Fig. S1”, Garrett et al., 2022b) and summed daily averages that were above 5 °C for this period. We calculated APD for this same period by adding the total amount of daily precipitation (mm). Temperature and precipitation data were retrieved at a spatial resolution of 50 x 50 km using the nasapower package (Sparks, 2024) ran in R (v.4.3.1; R core Team, 2023), which provides hourly climate data from the Prediction of Worldwide Energy Resource project.

Temperature and precipitation (and incidentally, nest humidity) can affect the activity, development, and survival of nestlings and their prey and parasites via both direct and indirect effects. Such effects encompass thermal stress on nestlings (de Zwaan et al. 2020) and their parents (Tapper et al. 2020), as well as reduced food-provisioning and insect prey availability under extreme weather conditions, typically on rainy cold snaps but also on heatwave days (Tapper et al. 2020, Garrett et al. 2022a). Effects of meteorological conditions may also extend to *Protocalliphora* parasitic loads (Robledo 2019, Castaño-Vázquez et al. 2021) by exerting a direct effect on the activity, development, and survival of *Protocalliphora* of all stages (Dawson et al. 2005, Mennerat et al. 2021), as well as through a reduction in nestling host quality and availability. We accordingly averaged the hourly temperature and precipitation data for each swallow brood during both the specific developmental period of its nestlings and *Protocalliphora* therein (see previous section).

### Statistical analyses

The causal pathways through which the habitat composition of agricultural landscapes, weather conditions, and the age of breeding females may affect the probability and intensity of Tree Swallow brood infestation by *Protocalliphora* likely include their effect on the hatching date and nestling availability of swallow broods as shown by the directed acyclic graph (DAG) in Figure 1. To assess the direct and indirect effects shown in this DAG, we had to fit four models (A-D), which response variables included the date of the first hatching event of a swallow brood (A), the number of nestling days that a swallow brood was available to *Protocalliphora* eggs and larvae (B), and the presence (C) and abundance (D) of *Protocalliphora* puparia in a swallow nest (Table 1). We determined the explanatory variables to include and control for in these models following Cinelli’s et al. (2022) recommendations, notably to avoid biased estimates caused by back-door paths. For instance, we had to include farm longitude in Model A because climatic conditions and agriculture intensity follow the East-West gradient in topography of our study area (Fig. 2A). Only broods containing *Protocalliphora* were retained in model D, as the effect size of factors driving the abundance of *Protocalliphora* in a brood (model D) could differ from those driving their occurrence (model C).

We estimated effect sizes by fitting Generalized Linear Mixed Models (GLMMs) or Generalized Additive Mixed Models (GAMMs) based on the nature of the response variables and the expected shape of their relationships with explanatory variables (i.e., strictly monotonous or not; Table 1). These models included farm IDs nested within years as random effects on the intercept to account for the hierarchical structure of the sampling scheme. We fitted GLMMs and GAMMs (following model G of Pedersen et al. 2019) using the glmmTMB (v.1.1.9; Brooks et al. 2017) or mgcv (v.1.8.42; Wood 2023) R packages, respectively. We assessed model assumptions with the *DHARMa* (v.0.4.7; Hartig 2022) R package, and generated model predictions at the population-level by marginalizing over the random effect structure and setting non-focal explanatory variables at their mean using the *marginaleffects* R package (v.0.18.0, Arel-Bundock, 2024). All explanatory variables were standardized to zero mean and unit variance except Comp.1 and Comp.2, the scores of the first two axes of the rPCA used to define agricultural landscape contexts. The conditional and marginal *R*² of the GLMM (model A) were calculated following Nakagawa’s et al. (2017) with the *MuMIn* package (v.1.47.5, Barton 2023). Given that random effects are treated as smooth term, which complicates variance partitioning in GAMMs (model B,C,D), we only provide the conditional *R*² returned by the *mgcv* summary function (Wood, 2023).

## Results

### Tree Swallow Hatching Phenology (Model A)

Hatching of the first egg ranged from DOY 146 to 194 across the 2,560 Tree Swallow clutches we monitored between 2004 and 2019, with a mean DOY (± SD) of 160.50 ± 6.69. The distribution of hatching dates was strongly skewed to the right and hatching events occurring past DOY 175 likely originated from second breeding attempts by females that experienced an early failure on their first attempt in another nest box. While the fixed effects we considered (Table 1, Fig. 1) explained 15.75% of the variance (marginal *R*^2^) in hatching date, this proportion increased to 30.98% with the addition of random effects (conditional *R*^2^), highlighting the importance of some environmental conditions that varied among farms and years that were not included in the analysis.

Eggs of ASY female swallows hatched 6 days (± 0.5 SE; 95% CI: 159.4; 161.0) sooner on average than those of younger SY females (*P* < 0.001; Fig. 3a). Spring weather (DOY 91-152) also affected the timing of first hatching events. While increased accumulated spring precipitation (APD) delayed hatching (*P* < 0.001; Fig. 3B), higher accumulated spring temperature (ADD) led to earlier hatching (*P* = 0.06; Fig. 3C). Agricultural context influenced the timing of hatching events only through Comp.1 with hatching events being delayed in more agrointensive areas (*P* < 0.001, Fig. 3D). Longitude had no direct effect on the date of the first hatching event (*P* = 0.10) once the mediating effects of spring weather and agricultural context were taken into account.

**Figure 3.**
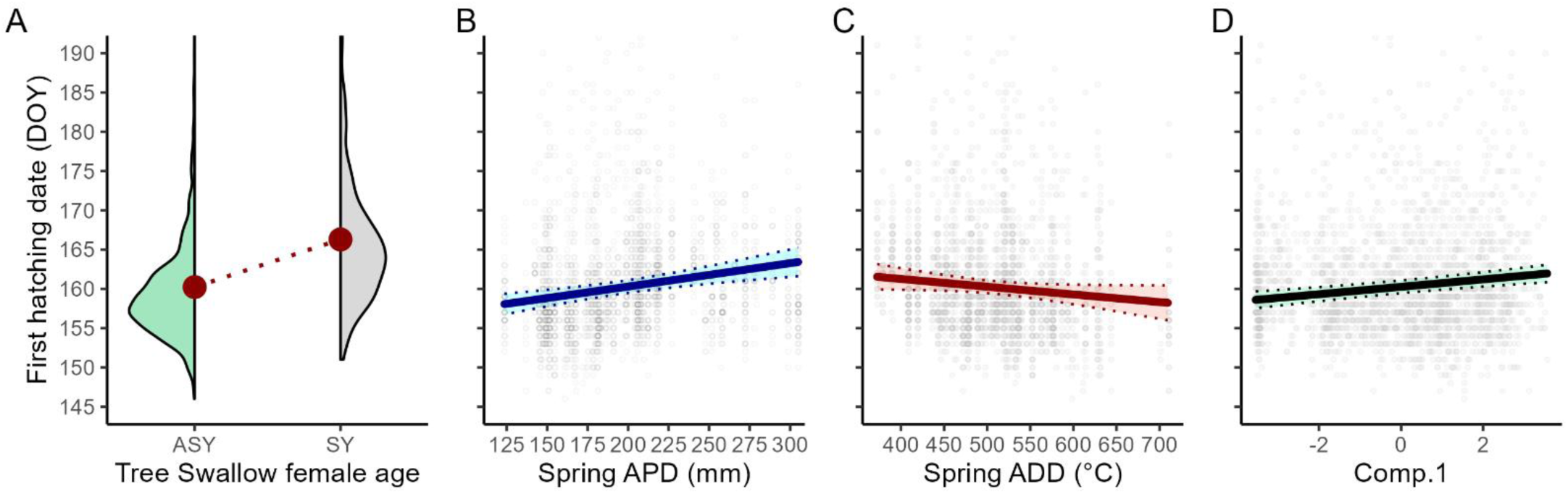
Predicted date (day of year or DOY) of the first hatching event in Tree Swallow broods according to A) the female swallow age as either After Second Year (ASY) or Second Year (SY); B) the spring Accumulated Precipitation Days (APD; mm); C) the spring Accumulated Degree Days (ADD; °C); and D) the agricultural context in which the swallow brood was raised (based on rPCA scores along Comp.1; see Fig. 2B). Predictions and 95% confidence interval were computed on the link-function scale while setting non-focal, fixed effects to their mean (or ASY for female age) and marginalizing over the random effect structure (see Table 1 for model details). Density kernels (A) and grey dots (B-D) depict raw data.

### Nestling Host Availability (Model B)

Nestling availability for *Protocalliphora* varied substantially among Tree Swallow broods with an average (± SD) of 93.9 ± 37.4 nestling days (Fig. 4A). The fixed and random effects of Model B (Table 1, Fig. 1) explained 32.3% of this variation.

**Figure 4.**
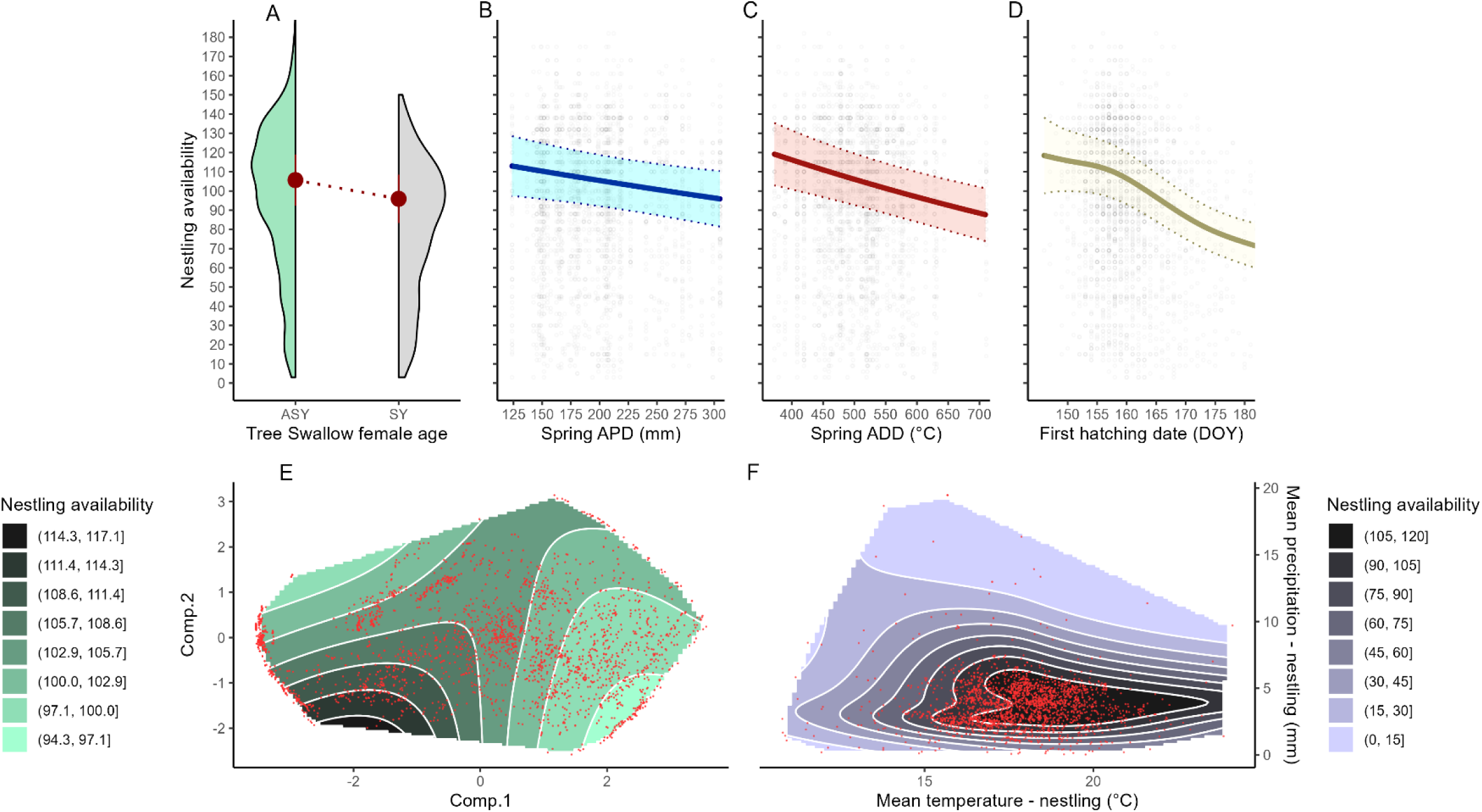
Predicted nestling host availability (nestling-days) provided by Tree Swallow broods to their *Protocalliphora* ectoparasites according to A) the female swallow age as either After Second Year (ASY) or Second Year (SY); B) the spring Accumulated Precipitation Days (APD); C) the spring Accumulated Degree Days (ADD); D) the date of the first hatching event of the swallow brood; E) the agricultural context in which the swallow brood was raised (based on rPCA scores along Comp.1 and Comp.2; see Fig. 2B); and F) the hourly mean temperatures and precipitation experienced by nestlings of a brood from the first hatching event to the last nestling’s fledging or death. Predictions and 95% confidence interval were computed on the link-function scale while setting non-focal, fixed effects to their mean (or ASY for female age) and marginalizing over the random effect structure (see Table 1 for model details). Density kernels (A) and grey or red dots (B-F) depict raw data.

The availability of nestlings provided by broods decreased at an increasing rate with their hatching date (*P* < 0.001; Fig. 4D). Beyond their effect on hatching date (Model A), the age of female Tree Swallows and spring weather (DOY 91-152) had a direct effect on nestling availability. Tree Swallow broods from ASY females provided a greater nestling availability to *Protocalliphora* than those of SY females (105.7 vs. 95.9 nestling days; *P* < 0.001; Fig. 4A). Nestling availability decreased linearly with both spring accumulated precipitation (APD; *P* = 0.02; Fig. 4B) and temperature (ADD; *P* < 0.001; Fig. 4C). That said, meteorological conditions during the nestling developmental period had by far the strongest impact on nestling availability with the interaction between the mean hourly precipitation and temperature defining a clear bivariate optimum (*P* < 0.001; Fig. 4F). Nestling availability peaked at a mean hourly precipitation and temperature of ∼4mm and ∼20 °C, respectively. Whereas nestling availability decreased sharply either below or above the precipitation optimum, it responded negatively to temperature mostly under cold conditions (< 17 °C).

Beyond its effect on hatching date (Model A), agricultural context, as defined by Comp.1, Comp.2 and their interaction, occasioned a direct saddle effect on nestling availability (*P* < 0.001; Fig. 4E). Controlling for the mediating effect of hatching date, nestling availability increased with the relative cover in forage and pastures as long as forest cover remained abundant in the landscape. It decreased, however, in forest dominated landscapes and reached its lowest in areas dedicated to row cropping and devoid of forest cover (Fig. 4E).

### Prevalence of *Protocalliphora* infestations (model C)

*Protocalliphora* have been found to infest 45.9% of the 2,560 Tree Swallow broods we sampled between 2004 and 2019. The likelihood of brood infestation, however, varies substantially among farms and years (Coroller-Chouraki et al. 2026). Model C (Table 1) addressed this variation and its fixed and random effects explained 26.1% of the variation in infestation probability.

Whereas hatching phenology had a limited, positive direct effect on infestation probability (*P* = 0.11; Fig. 5A), nestling availability had a strong and positive one (*P* < 0.001; Fig. 5B). Agricultural landscape context also had a substantial direct effect on infestation probabilities which varied from 0% to 88% under otherwise average conditions (*P* < 0.001; Fig. 5C). The highest infestation probabilities were associated with low Comp.1 and high Comp.2 values, indicative of landscapes with dense tree cover interspersed by pastures and hay fields. Conversely, infestation probability decreased with Comp.1 values, especially at high Comp.2 values, reaching its lowest in agrointensive landscapes dominated by corn and soybean crops. Spring meteorological conditions had no direct effect on infestation probability once controlling for the mediating effect of hatching date and nestling availability (*P* = 0.98 and *P* = 0.45 for cumulated precipitation and temperature, respectively). However, mean temperature and precipitation during *Protocalliphora* larval and pupation period implied infestation probability variations from 9% to 75% under otherwise average conditions (*P* = 0.001; Fig. 5D). *Protocalliphora* infestation probability increased with mean temperature and precipitation, yet the effect of temperature decreased as precipitation levels increased.

**Figure 5.**
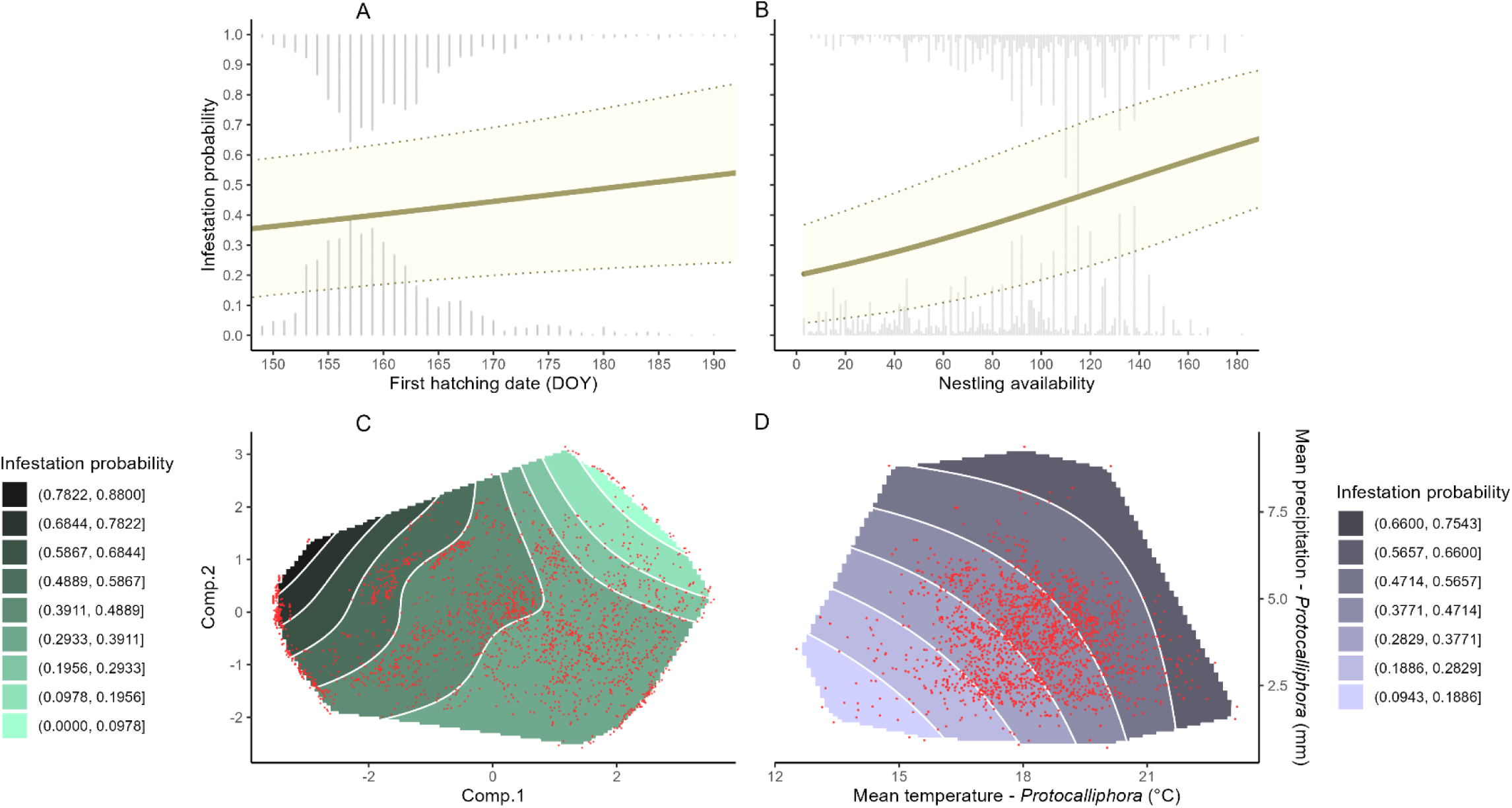
Predicted probability (prevalence) of Tree Swallow brood infestation by *Protocalliphora* blowflies according to A) the date of the first hatching event of the swallow brood; B) the nestling host availability for *Protocalliphora*; C) the agricultural context in which the swallow brood was raised (based on rPCA scores along Comp.1 and Comp.2; see Fig. 2B); and D) the hourly mean temperatures and precipitation experienced by *Protocalliphora* larvae and pupae. Predictions and 95% confidence interval were computed on the link-function scale while setting non-focal, fixed effects to their mean and marginalizing over the random effect structure (see Table 1 for model details). Grey bars (A-B) depict densities of presence (1) or absence (0) and red dots (C-D) depict raw data.

### *Protocalliphora* abundance within infested swallow broods (model D)

Overall, the count distribution of *Protocalliphora* puparia within infested Tree Swallow broods (*n* = 1,175) was rightly skewed with a maximum of 127 and a mean of 18.4 ± 18.6 (SD) puparia. The fixed and random effects of the model of *Protocalliphora* abundance within infested broods (Table 1) explained 25.8% of the variation in this ectoparasitic load.

Hatching time had a moderate direct effect on *Protocalliphora* abundance (*P* = 0.001; Fig. 6A). Mean abundance peaked between DOY 165 and 170, plateauing at 18 individuals, before it decreased progressively in later clutches, reaching 10 individuals in clutches that hatched around DOY 190. Nestling availability had, however, a substantial direct impact on *Protocalliphora* abundance (*P* < 0.001; Fig. 6B). Under otherwise average conditions, broods most likely to fail (i.e., with a nestling availability ≤ 20 nestling-days) had a mean abundance lower than 6 *Protocalliphora*, while broods with a nestling availability greater than 140 nestling-days harbored a mean abundance above 25 *Protocalliphora*. The direct influence of landscape context on *Protocalliphora* abundance (*P* = 0.001; Fig. 6C) was consistent with the pattern observed for the probability of nest infestation (Fig. 5C), creating a mean difference of ∼20 *Protocalliphora* across the agricultural intensity gradient under otherwise average conditions. The highest mean abundances of *Protocalliphora* occurred in areas characterized by low Comp.1 and high Comp.2 values, indicative of dense tree cover punctuated by pastures and hay fields. Mean abundance decreased with increasing Comp.1 values, reaching its lowest values in agrointensive landscapes dominated by corn and soybean crops. Spring meteorological conditions had no direct bearing on *Protocalliphora* abundance after controlling for the mediating effect of hatching date and nestling availability (*P* = 0.40 and *P* = 0.62 for cumulated precipitation and temperature, respectively). Yet those occurring during *Protocalliphora* larval development and pupation had a strong direct effect on abundance (Fig. 6D; *P* = 0.010). Under otherwise average conditions, abundance increased at an increasing rate with both mean temperature and precipitation, causing a 5.5-fold increase in *Protocalliphora* mean abundance when moving from the coldest and driest conditions (∼6 puparia) to the warmest and wettest conditions (∼30 puparia), respectively.

**Figure 6.**
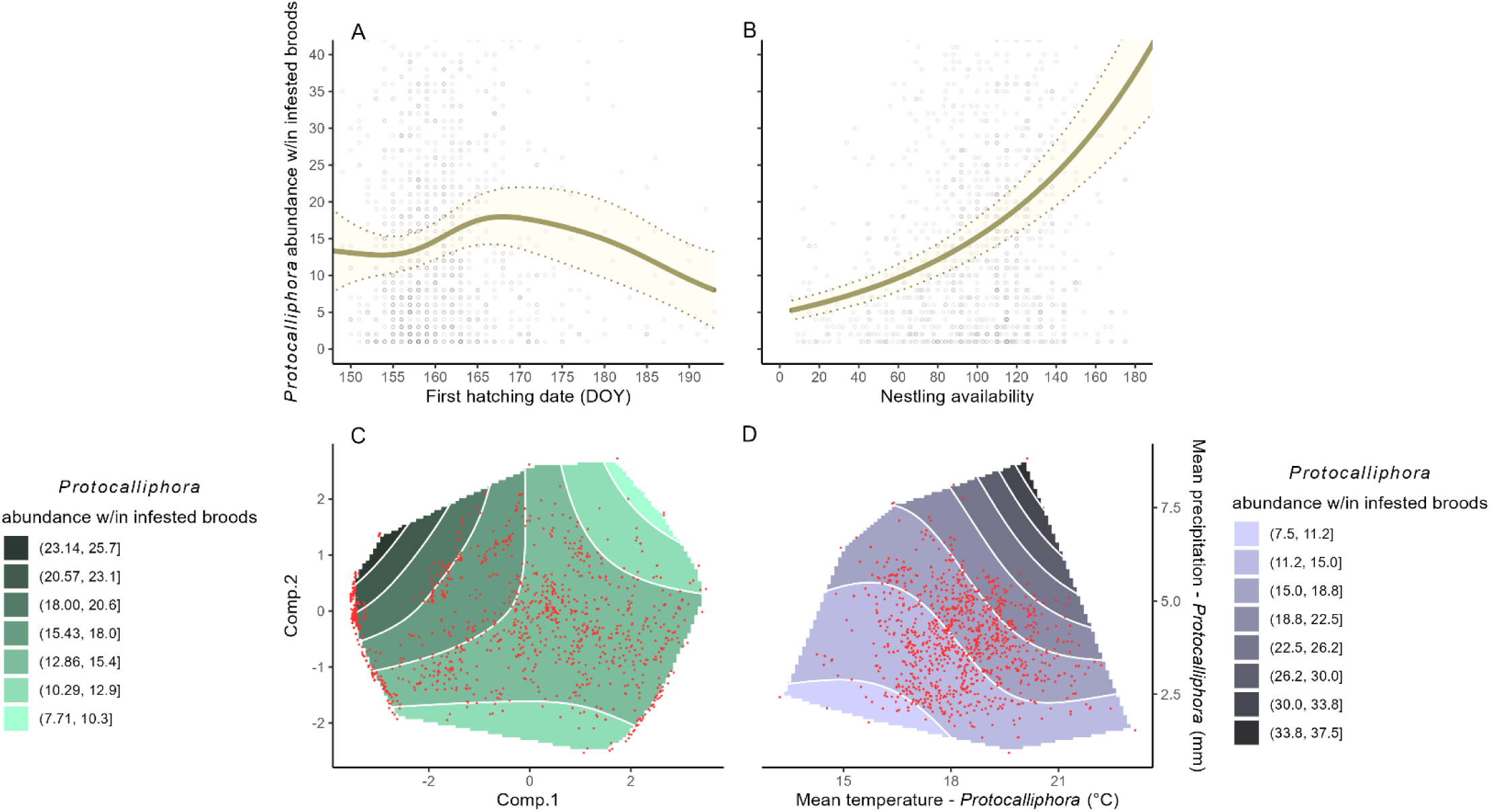
Predicted abundance (load) of *Protocalliphora* puparia within infested Tree Swallow broods according to A) the date of the first hatching event of the swallow brood; B) the nestling host availability for *Protocalliphora*; C) the agricultural context in which the swallow brood was raised (based on rPCA scores along Comp.1 and Comp.2; see Fig. 2B); and D) the hourly mean temperatures and precipitation experienced by *Protocalliphora* larvae and pupae. Predictions and 95% confidence interval were computed on the link-function scale while setting non-focal, fixed effects to their mean and marginalizing over the random effect structure (see Table 1 for model details). Grey and red dots depict raw data.

## Discussion

Given the intricate pathways by which species can interact, understanding the ecological consequences of agricultural intensification requires looking beyond its direct effects on a single species or functional guild and considering its broader impacts on ecological networks. For instance, most populations of farmland birds and their insect prey have been found to decline with agricultural intensification around the Globe (Stanton et al. 2018, Raven and Wagner 2021, Outhwaite et al. 2022, Rigal et al. 2023), suggesting the importance of trophic relationships to explain some of the impacts of this anthropogenic environmental transformation (Boatman et al. 2004, Morris et al. 2005, Poulin et al. 2010, Goebel et al. 2024)). While predation is certainly important in that respect, other types of interactions such as parasitism must also be considered to understand how agriculture can affect species and their interactions, and how these effects may or may not propagate among species.

Using a 16-year dataset (2004-2019) comprising 2,560 Tree Swallow broods, we showed that agriculture intensity not only affected negatively the number and survival of nestlings, but also the prevalence and loads of *Protocalliphora* blowflies parasitizing broods. Notably, agriculture intensity reduced the likelihood and magnitude of *Protocalliphora* infestations both directly (through mechanisms yet to be quantified) and indirectly by limiting host availability for blowflies within swallow nests. This takes on particular significance when considering the negative effect of agriculture intensity on the number and body condition of fledglings produced by farmland birds, including Tree Swallows, and this as found by studies that had already controlled for its negative impact on the birds’ food resources (e.g., Benton et al. 2002, Garrett et al. 2022b). If agriculture intensity indeed reduces the parasitic pressure exerted by *Protocalliphora* blowflies on nestlings, and thereby their impact on fledglings’ body condition and eventual recruitment to the breeding population (Thomas et al. 2007), this would suggest that agricultural impacts beyond food limitation and parasitism, such as toxicological effects resulting from pesticide use (e.g., (Stanton et al. 2018, Poisson et al. 2021, Molenaar et al. 2024), are more important than previously recognized. Our work highlights the need to assess the impacts of ecological perturbations through an integrative, systemic, and causal approach to mitigate biases and prevent misinterpretations of the putative roles of multiple, interacting stressors.

### Swallow breeding phenology

In migratory birds, individuals that arrive earlier on breeding grounds are typically of higher quality, often being older (more experienced) or in better physical condition than those arriving later. Consequently, they tend to secure higher quality nesting sites (Verhulst and Nilsson 2007, Sergio et al. 2009, Sumasgutner et al. 2019), unless mistaken by ecological traps (Hale and Swearer 2016, Courtois et al. 2021). We found that maternal age was a key predictor of hatching phenology in Tree Swallows, with older and more experienced females producing nestlings that hatched ∼6 days earlier than those of second-year mothers. A pattern consistent across most North-American nest-box networks where Tree Swallow breeding activities are monitored (Robertson and Rendell 2001, Winkler et al. 2020, Courtois et al. 2021).

Hatching of Tree Swallow clutches occurred earlier with warmer temperatures from April to June and later with higher rainfall during that same period. These patterns concord with those found for Tree Swallows and several other passerine bird species inhabiting temperate regions (e.g., Bourret et al. 2015, Imlay et al. 2019, de Zwaan et al. 2022, Taff and Shipley 2023). Spring weather at the breeding site thus appears to be a strong determinant of Tree Swallow breeding phenology, as observed in many other long-distance migrants (Ockendon et al. 2013). Spring weather likely exert an indirect influence on avian breeding phenology through insect emergence and trophic availability for birds, *inter alia* through green-up phenology (Burke 2014, Thorup et al. 2017, Gow et al. 2019, Belitz et al. 2025). Our observation of delayed hatching in agrointensive landscape contexts may also result from a mediating effect of insect trophic availability. Indeed, trophic availability for Tree Swallows was shown to be lower in agrointensive landscape contexts of our study area at the beginning of the breeding period (Garrett et al. 2022c). In such habitats, female Tree Swallows may require an extended period to forage and accumulate reserves before initiating energetically demanding breeding activities (Gillies et al. 2022), or may need to more closely synchronize peak nestling food demands with periods of higher resource availability (Paquette et al. 2013). Hence, female age, spring weather and agroenvironmental conditions can all contribute to shape avian breeding phenology, with potential influences on the degree of synchrony between birds and insect communities at breeding sites (Renner and Zohner, 2018) and thus implications for both insect trophic availability for nestlings and their exposure to insect ectoparasites.

### Nestling host availability for *Protocalliphora*

Partly correlated with breeding success, the number of nestling-days reflects the level and stability of blood availability and temperature generated by nestlings that can be exploited by *Protocalliphora* blowfly larvae. We found that it increased with Tree Swallow maternal age and decreased with hatching date as observed across a wide array of bird species and guilds (e.g., (Ghilain and Bélisle 2008, Winkler et al. 2013, Shipley et al. 2020, Garrett et al. 2022a). While earlier breeding is typically associated with larger clutches and higher nestling survival, it also carries the potential thermoregulatory and dietary costs of variable and extreme weather conditions and lower insect prey availability that can be harmful if not lethal to nestlings (Winkler et al. 2013, Shipley et al. 2020, Garrett et al. 2022a,c). Nestling availability for *Protocalliphora* peaked between mean hourly temperatures of 17.0 °C and 23.0 °C and when hourly precipitations averaged 2.5mm to 6.0mm during the nestling development. Importantly, nestling availability declined sharply when temperatures dropped below 17.0 °C, which coincides with the significant reduction in insect prey availability for Tree Swallows at a threshold 18.2 °C observed in our study area (Garrett et al. 2022a). This result not only underscores the importance of integrating local meteorological data to address direct and indirect effects on the reproductive success of insectivorous birds (Weegman et al. 2017) but also on their insect ectoparasites.

Nestling availability was higher in forage-dominated landscapes, particularly those with low tree cover, and lower in agrointensive landscapes. This pattern likely arises because survival, growth, and fledging success of Tree Swallow nestlings in our study area exhibit consistent trends that are sufficiently strong to compensate for the opposite trend observed in nestling period duration (Garrett et al. 2022a,b). These patterns have in turn be attributed to the negative effects of agriculture intensity on Tree Swallow insect trophic availability (Bellavance et al. 2018) and food provisioning rate (Garrett et al. 2022b). While the contamination of Tree Swallow insect prey by pesticides also increases with the studied agriculture intensity gradient, no clear direct (toxicological) or indirect (trophic) effect of these contaminants on either nestlings or adults have so far been found (Poisson et al. 2021, Sigouin et al. 2021). Recent meta-analyses suggest that neonicotinoid insecticides not only reduce insect prey availability but also disrupt bird behavior and survival, with significant negative effects on their population dynamics (Molenaar et al. 2024). These investigations, however, did not adopt an integrative, causal framework that would allow assessment of the relative importance of the different pathways through which pesticides may affect the various ecological components of farmland bird breeding success, including the likelihood and severity of ectoparasite infestations and their consequences for these birds.

### *Protocalliphora* prevalence and load in agroenvironmental contexts

Nestling-days emerged as a strong determinant of both *Protocalliphora* prevalence and load in Tree Swallow broods. Although this study is the first to use nestling-days as a proxy for host availability, our results are consistent with previous work generally reporting a positive effect of brood size on *Protocalliphora* and other ectoparasite loads in passerines (e.g., Hurtrez-Boussès et al. 1999, Tomás et al. 2007, Dawson et al., 2011; but see Remeš and Krist, 2005). Larger and longer-lasting clutches and broods likely provide stronger visual and olfactory cues for *Protocalliphora* adults to locate nests and oviposit (Williams 2017). In addition, prolonged nestling availability ensures sustained access to blood meals and a nest microhabitat potentially favorable to ectoparasite development (Albert et al. 2023). Although high *Protocalliphora* loads have been hypothesized to impair host breeding success (by slowing nestling growth, delaying fledging, or reducing fledging mass and success; Johnson and Albrecht 1993, Simon et al. 2004, Hannam 2006), their effects may be highly variable among host-parasite systems, geographic regions, and environmental contexts (Bennett and Whitworth 1992, Newton 1998, Musgrave et al., 2019). When detected, impacts on fledging mass or success have been shown to depend upon environmental conditions, particularly weather and nestling food availability (Howe 1992, Merino and Potti 1996, Musgrave et al. 2019). While *Protocalliphora* parasitism likely adds to or exacerbates other stressors experienced by nestlings, its effects are more likely to manifest itself in terms of body condition (e.g., hematocrit and immune function) and post-fledging survival (Thomas et al. 2007, Poisson et al. 2021). Consequently, and consistent with the assumption of our DAG (Fig. 1), the causal effect of nestling availability (as indexed by nestling-days) on *Protocalliphora* load is likely stronger than the reverse.

Although weather conditions play a key role in shaping the phenology and growth conditions of passerines (Gow et al. 2019, Imlay et al. 2019, de Zwaan et al. 2020), as well as those of their insect prey and parasites (Dawson et al. 2005, Forrest 2016, Renner and Zohner 2018), interpreting and comparing their effects on the prevalence and severity of *Protocalliphora* infestations remains challenging. This difficulty arises largely because existing studies differ substantially in their biological systems, environmental contexts, and analytical approach. For example, studies have focused on different host-parasite species (e.g., migratory vs. resident or open-cup vs. cavity nesting birds; (Mason 1944, Bennett and Whitworth 1992, Heeb et al. 2000), have been conducted under contrasting climatic regimes (e.g., Mediterranean in Corsica vs. temperate, humid continental in South-Eastern Canada; (Bańbura et al. 2004, Hannam 2006), and have considered weather variables at different temporal resolution and over different time windows (e.g., (Musgrave et al., 2019, Mennerat et al. 2021). In addition, using model selection methods, studies have variably controlled (or failed to adequately control; *sensu* Cinelli et al. 2022) for potential confounding or mediating factors associated to weather, such as laying or hatching date and availability of nestlings within nests (Mennerat et al. 2021, Maziarz et al. 2022). This is particularly important because, as implied by our DAG (Fig. 1) and supported by our results (Fig. 5 and 6), hatching date is likely to influence *Protocalliphora* prevalence and load primarily through its indirect effect on weather conditions (Section S2; Figure S2) experienced by nestlings and their ectoparasites, resulting in a weak direct effect on infestation metrics. Adopting a causal inference framework can help limit such biases, provided that the working DAG accurately reflects the data-generating process (Cinelli et al. 2022). On that front, no study to date (including ours) has so far accounted for the fact that bird blowfly communities may comprise multiple species, many of which parasitize a broad range of bird host (Bennett and Whitworth 1992). Despite limited knowledge of the host selection and oviposition behavior of bird blowflies, it is reasonable to hypothesize that species-specific differences in bird host phenology and abundance may generate temporal variation in nest availability, and thereby influence *Protocalliphora* infestation prevalence and severity. Future studies would therefore benefit from adopting a more integrative community-level approach that explicitly accounts for host and parasite diversity and their phenological dynamics.

Most notably, this work extends and strengthens the 2-year exploratory study of Daoust et al. (2012a) and represent the first long-term investigation of how agricultural intensity modulates host-parasitism relationships between a wild bird species and its primary insect ectoparasites. Both the prevalence and intensity of *Protocalliphora* infestations in Tree Swallow nests were higher in landscapes dominated by pastures and forage crops embedded within extensive forest cover. In contrast, prevalence and intensity were lower in agrointensive landscapes lacking forest cover and dominated by annual row crops, primarily maize, soybean and other cereals. Although nestling availability within infested nests emerged as a key determinant of *Protocalliphora* prevalence and load (Fig. 5 and 6), it showed only a moderate decline along the agricultural intensity gradient (Fig. 4). This pattern suggests that agricultural practices within 500 m of swallow nests are unlikely to have substantially influenced *Protocalliphora* infestations through their effects on nestling availability in Tree Swallow nests. Multiple causal pathways could explain the (direct) influence of agricultural intensity on *Protocalliphora* infestation prevalence and intensity.

First, agrointensive landscapes may be less favorable to adult *Protocalliphora* prior to oviposition because they provide fewer food resources (i.e., nectar, flowers, and fruits; Bennett and Whitworth, 1991) and less shelter, particularly for overwintering (Bennett and Whitworth 1992, Daoust et al. 2012a). Second, many *Protocalliphora* species, including those examined here (i.e., *P. sialia, P. bennetti,* and *P. metallica*; (Coroller-Chouraki et al. 2026) are capable of infesting the nests of multiple bird species (Bennett and Whitworth 1992). Agrointensive landscapes in our study area (Robert et al. 2019, Le Vaillant et al. 2026) as in much of North America and Europe (Stanton et al. 2018, Rigal et al. 2023), support lower avian species and abundance, generally accompanied by reduced breeding success, and may therefore provide lower host availabilities to bird blowflies in terms of both nest and nestling densities. By contrast, environments supporting more diverse and denser avian host communities may promote spillover processes (Daszak et al. 2000), whereby larger local pools of potential hosts facilitate host-parasite association, transmission, and persistence (Wells et al. 2015), ultimately increasing local *Protocalliphora* densities. Third, Tree Swallows and their dipteran prey are exposed to higher numbers and concentration of pesticides in agrointensive landscapes of our study area (Poisson et al. 2021), a situation that likely also affects their *Protocalliphora* ectoparasites (Sigouin et al. 2021). Beyond their lethal effects, many pesticides such as neonicotinoids (which were ubiquitous in our study area; Montiel-León et al. 2019, Giroux, 2022), are known to alter multiple aspects of insect biology, including phenology, behavior, and life-history traits (Pisa et al. 2015). Consequently, *Protocalliphora* inhabiting agrointensive landscapes may exhibit impaired host-searching and oviposition behaviors, reduced offspring quality, or increased mortality of larvae prior to pupation. These effects, however, may also extend to *Nasonia* parasitoid wasps (Hymenoptera: Pteromalidae), which are likely contribute to the regulation of *Protocalliphora* populations (Daoust et al. 2012a, Schöfer et al. 2023). The relative importance of these alternative, yet non-mutually exclusive, mechanistic pathways will benefit from a more integrative, community-level approach.

Although this study considers landscapes dominated by pastures and forage crops as representing the agroextensive end of the gradient used to assess the effects of agricultural intensification on interactions between Tree Swallows and their *Protocalliphora* ectoparasites, this classification requires nuance. Indeed, research in Europe has shown that pesticides (i.e., fungicides and insecticides) and anti-parasitic substances used in dairy and cattle farming can enter pastures and hayfields via livestock excreta or manure application, with detrimental consequences for the entomofauna (Mahdjoub et al. 2020, Buijs et al. 2022). In addition, improved growing conditions due to climate change, increasingly efficient harvest methods, and the widespread adoption of fast-growing crop varieties have resulted in earlier, faster, and more frequent forage harvest within a season, with negative effects on both insects and ground-nesting farmland birds (McGowan et al. 2021). Together, these factors suggest that, despite the strong negative effects of agrointensive contexts on *Protocalliphora* infestations of Tree Swallow nests relative to agroextensive ones, *Protocalliphora* communities may already be experiencing pesticide- and management-related declines in most modern farmlands.

This study provides the first long-term investigation of avian ectoparasitic infestation dynamics within agroecosystems, while also highlighting key avenues for future research. From a farmland bird conservation perspective, it remains to be determined whether the reduced prevalence and load of *Protocalliphora* in bird broods raised in agrointensive landscapes offset the multiple other fitness costs imposed by agricultural intensification. Beyond effects on parasite prevalence and load, agricultural intensification may affect *Protocalliphora* life-history traits, including body size and fecundity (Verheyen and Stoks 2019, Gérard et al. 2022, Rawal et al. 2025) as well as pupation success (Heneberg et al. 2020, Mahdjoub et al. 2020). Of particular interest are the potential effects of agricultural intensification on the development and emergence of *Protocalliphora* pupae, as successful emergence is critical for blowfly population persistence across breeding seasons. Although empirical data remain scarce, agricultural intensification may alter these processes, notably by modifying interactions with *Nasonia* parasitoid wasps (Daoust et al. 2012a, Haan et al. 2020, Coroller-Chouraki et al. 2026). Together, these considerations emphasize the need for integrative approaches to disentangle the cascading effects of agricultural intensification on host-parasite-parasitoid networks. In this respect, blowfly puparia remaining within avian nest material may represent valuable biological archives and potential bio-indicators for assessing the impacts of agricultural intensification impacts on non-target trophic networks within agroecosystems.

## Acknowledgments

We sincerely thank the 40 farm owners who generously agreed to participate in this long-term study initiated in 2004. We are also grateful to the many graduate students, as well as field and laboratory assistants, who contributed over the years to data collection, data entry, analyses and verification, for both avian and insect samples. Special thanks to Marianne Cusson for the illustrations included in the DAG. This research was carried out with the approval of the Université de Sherbrooke Animal Care Committee. Financial support was provided by Discovery Grants from the Natural Sciences and Engineering Research Council of Canada awarded to Fanie Pelletier, Dany Garant and Marc Bélisle; research grants from the Fonds de recherche du Québec - Nature et technologies to FP, DG and MB; the Canada Research Chairs program to FP and MB; the Canada Foundation for Innovation to FP, DG, and MB; the Canadian Wildlife Service of Environment and Climate Change Canada, and the Université de Sherbrooke.

