## Appendix 1 for "Enemy mitigation in farmlands: Intensive agriculture impacts avian nesting and subsequent ectoparasite infestation"

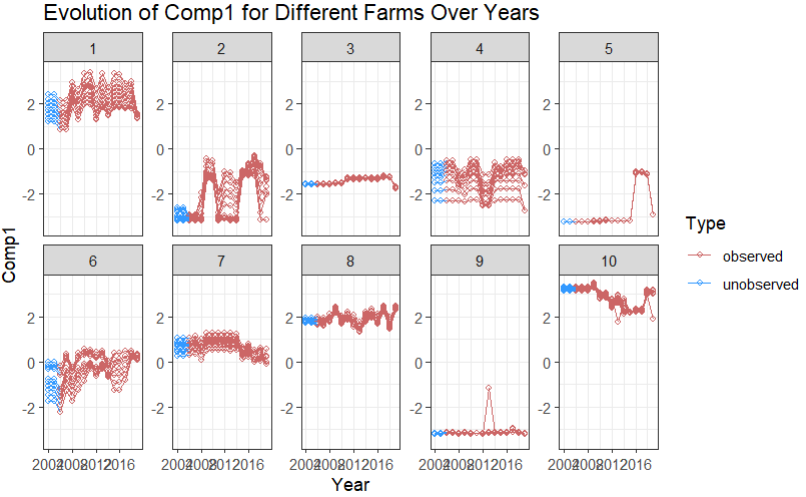

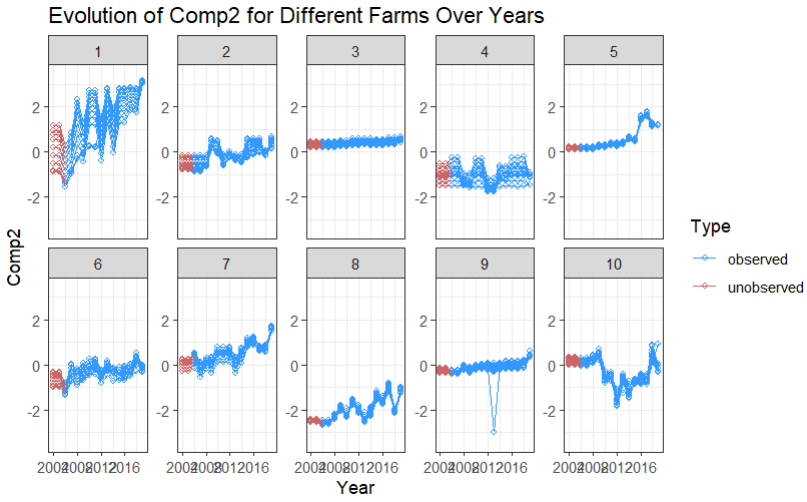

**Figure S1.** *Interannual stability of landscape composition around Tree Swallow nest boxes between 2004 and 2019. Extrapolated Comp.1 or Comp.2 for 2004 and 2005 missing values are displayed in contrasting color (respectively blue vs red, and red vs blue).*

**Section S1.** *Model outputs corresponding to A) First hatching date, B) Nestling availability in a nest, C) Infestation probability, D) Protocalliphora abundance w/in infested broods.*

- ***Tree Swallow Hatching Phenology (Model A)***

```
summary(Model_timing)
Family: Gamma  ( log )
Formula:
Timing_clutch ~ (1 | Year/Farm) + scl_A1J1_T + scl_A1J1_P +    scl_Longitude + Comp1 * Comp2 + Agemom
Data: data_ch2
Random effects:
Conditional model:
 Groups    Name        Variance  Std.Dev.
 Farm:Year (Intercept) 1.599e-04 0.01264
 Year      (Intercept) 9.254e-05 0.00962
Number of obs: 2560, groups:  Farm:Year, 583; Year, 16
Dispersion estimate for Gamma family (sigma^2): 0.00114
Conditional model:
                Estimate Std. Error z value Pr(>|z|)
(Intercept)    5.0730021  0.0025907  1958.2  < 2e-16 ***
scl_A1J1_T    -0.0040678  0.0021776    -1.9   0.0618 .
scl_A1J1_P     0.0081396  0.0020562     4.0 7.54e-05 ***
scl_Longitude  0.0028090  0.0016850     1.7   0.0955 .
Comp1          0.0029833  0.0006584     4.5 5.86e-06 ***
Comp2         -0.0010064  0.0009510    -1.1   0.2899
AgemomSY       0.0371484  0.0020270    18.3  < 2e-16 ***
Comp1:Comp2    0.0002321  0.0005482     0.4   0.6720
---
Signif. codes:  0 '***' 0.001 '**' 0.01 '*' 0.05 '.' 0.1 ' ' 1
```

- ***Tree Swallow Nestling Availability (Model B)***

```
summary(Model_chicks)
Family: gaussian
Link function: log
Formula:
cumulated_chicks ~ s(Farm, by = Year, bs = "re") + s(Year, bs = "re") + Agemom + scl_A1J1_P + scl_A1J1_T + s(scl_Timing_clutch) + te(Comp1, Comp2) + te(scl_mean_T_bird, scl_mean_P_bird)
Parametric coefficients:
            Estimate Std. Error t value Pr(>|t|)
(Intercept)  4.51984    0.02182 207.150  < 2e-16 ***
AgemomSY    -0.09668    0.02226  -4.343 1.47e-05 ***
scl_A1J1_P  -0.03991    0.01709  -2.335   0.0196 *
scl_A1J1_T  -0.06015    0.01449  -4.152 3.41e-05 ***
---
Signif. codes:  0 '***' 0.001 '**' 0.01 '*' 0.05 '.' 0.1 ' ' 1
Approximate significance of smooth terms:
                                          edf Ref.df      F  p-value
s(Farm):Year2004                     0.049155 40.000  0.001 0.496022
s(Farm):Year2005                     6.928911 39.000  0.224 0.147089
s(Farm):Year2006                     7.848384 38.000  0.294 0.067864 .
s(Farm):Year2007                     7.870476 36.000  0.298 0.096162 .
s(Farm):Year2008                     5.722161 36.000  0.199 0.173754
s(Farm):Year2009                     0.817394 29.000  0.029 0.403302
s(Farm):Year2010                     0.006617 39.000  0.000 0.772618
s(Farm):Year2011                     8.370360 37.000  0.400 0.012083 *
s(Farm):Year2012                    12.496630 33.000  0.717 0.005044 **
s(Farm):Year2013                     0.004668 36.000  0.000 0.916057
s(Farm):Year2014                     4.451686 38.000  0.145 0.181829
s(Farm):Year2015                    10.637277 39.000  0.422 0.031606 *
s(Farm):Year2016                     5.832156 33.000  0.231 0.157189
s(Farm):Year2017                     0.307489 36.000  0.009 0.431173
s(Farm):Year2018                     0.006801 38.000  0.000 0.765851
s(Farm):Year2019                     1.348776 36.000  0.040 0.364848
s(Year)                             12.051602 15.000  6.197  < 2e-16 ***
s(scl_Timing_clutch)                 3.426466  4.273 23.010  < 2e-16 ***
te(Comp1,Comp2)                      6.703794  8.009  3.521 0.000471 ***
te(scl_mean_T_bird,scl_mean_P_bird) 15.938783 17.732 18.674  < 2e-16 ***
---
Signif. codes:  0 '***' 0.001 '**' 0.01 '*' 0.05 '.' 0.1 ' ' 1
R-sq.(adj) =  0.323   Deviance explained = 35.3%
-REML =  12508  Scale est. = 945.16    n = 2560
```

- ***Protocalliphora presence within nests (model C)***

```
summary(presence_absence_protocalliphora)
Family: binomial
Link function: logit
Formula:
n_Protos ~ s(Farm, by = Year, bs = "re") + s(Year, bs = "re") +
scl_A1J1_P + scl_A1J1_T + s(scl_Timing_clutch) + scl_cumulated_chicks +            te(Comp1, Comp2) + te(scl_mean_T_protos, scl_mean_P_protos)
Parametric coefficients:
                      Estimate Std. Error z value Pr(>|z|)
(Intercept)          -0.213162   0.131989  -1.615    0.106
scl_A1J1_P            0.002879   0.117387   0.025    0.980
scl_A1J1_T           -0.079876   0.105516  -0.757    0.449
scl_cumulated_chicks  0.400417   0.053295   7.513 5.77e-14 ***
---
Signif. codes:  0 '***' 0.001 '**' 0.01 '*' 0.05 '.' 0.1 ' ' 1
Approximate significance of smooth terms:
                                           edf Ref.df Chi.sq p-value
s(Farm):Year2004                        18.930 40.000 37.937 0.00115 **
s(Farm):Year2005                        10.139 39.000 14.065 0.06781 .
s(Farm):Year2006                        17.704 38.000 33.553 0.00217 **
s(Farm):Year2007                        14.797 36.000 25.770 0.01217 *
s(Farm):Year2008                        13.573 36.000 21.306 0.03821 *
s(Farm):Year2009                         4.719 29.000  5.815 0.21167
s(Farm):Year2010                         1.757 39.000  1.818 0.40685
s(Farm):Year2011                        10.922 37.000 15.826 0.06410 .
s(Farm):Year2012                        11.891 33.000 18.996 0.03577 *
s(Farm):Year2013                         8.693 36.000 11.919 0.09973 .
s(Farm):Year2014                        16.935 38.000 33.524 0.00463 **
s(Farm):Year2015                        11.647 39.000 18.020 0.03103 *
s(Farm):Year2016                         6.670 33.000  8.630 0.15234
s(Farm):Year2017                        11.331 36.000 17.256 0.05047 .
s(Farm):Year2018                        18.352 38.000 33.829 0.00761 **
s(Farm):Year2019                        12.440 36.000 20.361 0.01785 *
s(Year)                                 10.826 15.000 99.540 2.9e-05 ***
s(scl_Timing_clutch)                     1.000  1.001  2.531 0.11174
te(Comp1,Comp2)                         10.098 12.259 89.884 < 2e-16 ***
te(scl_mean_T_protos,scl_mean_P_protos)  3.001  3.002 16.238 0.00102 **
---
Signif. codes:  0 '***' 0.001 '**' 0.01 '*' 0.05 '.' 0.1 ' ' 1
R-sq.(adj) =  0.261   Deviance explained = 26.1%
-REML = 1555.1  Scale est. = 1         n = 2560
```

- ***Protocalliphora abundance within infested nests (model D)***

```
summary(abondance_protocalliphora_without_0)
Family: Tweedie(p=1.99)
Link function: log
Formula:
n_Protos ~ s(Farm, by = Year, bs = "re") + s(Year, bs = "re") + scl_A1J1_T + scl_A1J1_P + s(scl_Timing_clutch) + scl_cumulated_chicks + te(Comp1, Comp2) + te(scl_mean_T_protos, scl_mean_P_protos)
Parametric coefficients:
                     Estimate Std. Error t value Pr(>|t|)
(Intercept)           2.70318    0.06378  42.382   <2e-16 ***
scl_A1J1_T           -0.02796    0.05585  -0.501    0.617
scl_A1J1_P           -0.05001    0.05823  -0.859    0.391
scl_cumulated_chicks  0.37255    0.02978  12.511   <2e-16 ***
---
Signif. codes:  0 '***' 0.001 '**' 0.01 '*' 0.05 '.' 0.1 ' ' 1
Approximate significance of smooth terms:
                                              edf Ref.df     F  p-value
s(Farm):Year2004                         7.389057 28.000 0.345  0.16437
s(Farm):Year2005                         0.004714 35.000 0.000  0.67485
s(Farm):Year2006                         9.377990 27.000 0.556  0.04764 *
s(Farm):Year2007                         0.004956 28.000 0.000  0.56505
s(Farm):Year2008                         0.001834 31.000 0.000  0.81254
s(Farm):Year2009                         0.001172 14.000 0.000  0.95000
s(Farm):Year2010                        10.896538 27.000 0.722  0.01423 *
s(Farm):Year2011                         6.921657 29.000 0.349  0.07724 .
s(Farm):Year2012                         7.562841 21.000 0.556  0.08885 .
s(Farm):Year2013                         1.988211 29.000 0.070  0.42491
s(Farm):Year2014                         0.002154 33.000 0.000  0.88515
s(Farm):Year2015                         0.001697 28.000 0.000  0.71863
s(Farm):Year2016                         4.253722 25.000 0.202  0.25420
s(Farm):Year2017                         7.806210 26.000 0.399  0.17549
s(Farm):Year2018                        12.428060 30.000 0.887  0.00179 **
s(Year)                                  8.813674 14.000 3.864 1.43e-05 ***
s(scl_Timing_clutch)                     3.751462  4.608 4.048  0.00261 **
te(Comp1,Comp2)                          6.166136  7.840 3.249  0.00136 **
te(scl_mean_T_protos,scl_mean_P_protos)  4.742882  5.573 2.699  0.01696 *
---
Signif. codes:  0 '***' 0.001 '**' 0.01 '*' 0.05 '.' 0.1 ' ' 1
R-sq.(adj) =  0.259   Deviance explained = 29.8%
-REML = 4243.4  Scale est. = 0.81326   n = 1108
```

**Section S2.** *Model output and prediction corresponding to the relation between Hatching Date (phenology) and mean temperature (°C) during nestling development.*

- ***Mean temperature during nestling development vs Hatching date (additionnal model)***

```
Family: gaussian  ( identity )
Formula:          scl_mean_T_bird ~ Timing_clutch + (1 | Year/Farm)
Data: data_ch2
      AIC       BIC    logLik -2*log(L)  df.resid
   5383.2    5412.5   -2686.6    5373.2      2555
Random effects:
Conditional model:
 Groups    Name        Variance Std.Dev.
 Farm:Year (Intercept) 0.2002   0.4474
 Farm:Year (Intercept) 0.2002   0.4474
 Year      (Intercept) 0.2006   0.4478
 Residual              0.3592   0.5993
Number of obs: 2560, groups:  Farm:Year, 583; Year, 16

Dispersion estimate for gaussian family (sigma^2): 0.359
Conditional model:
                Estimate Std. Error z value Pr(>|z|)
(Intercept)   -11.288090   0.360283  -31.33   <2e-16 ***
Timing_clutch   0.070167   0.002126   33.01   <2e-16 ***
---
Signif. codes:  0 '***' 0.001 '**' 0.01 '*' 0.05 '.' 0.1 ' ' 1
```

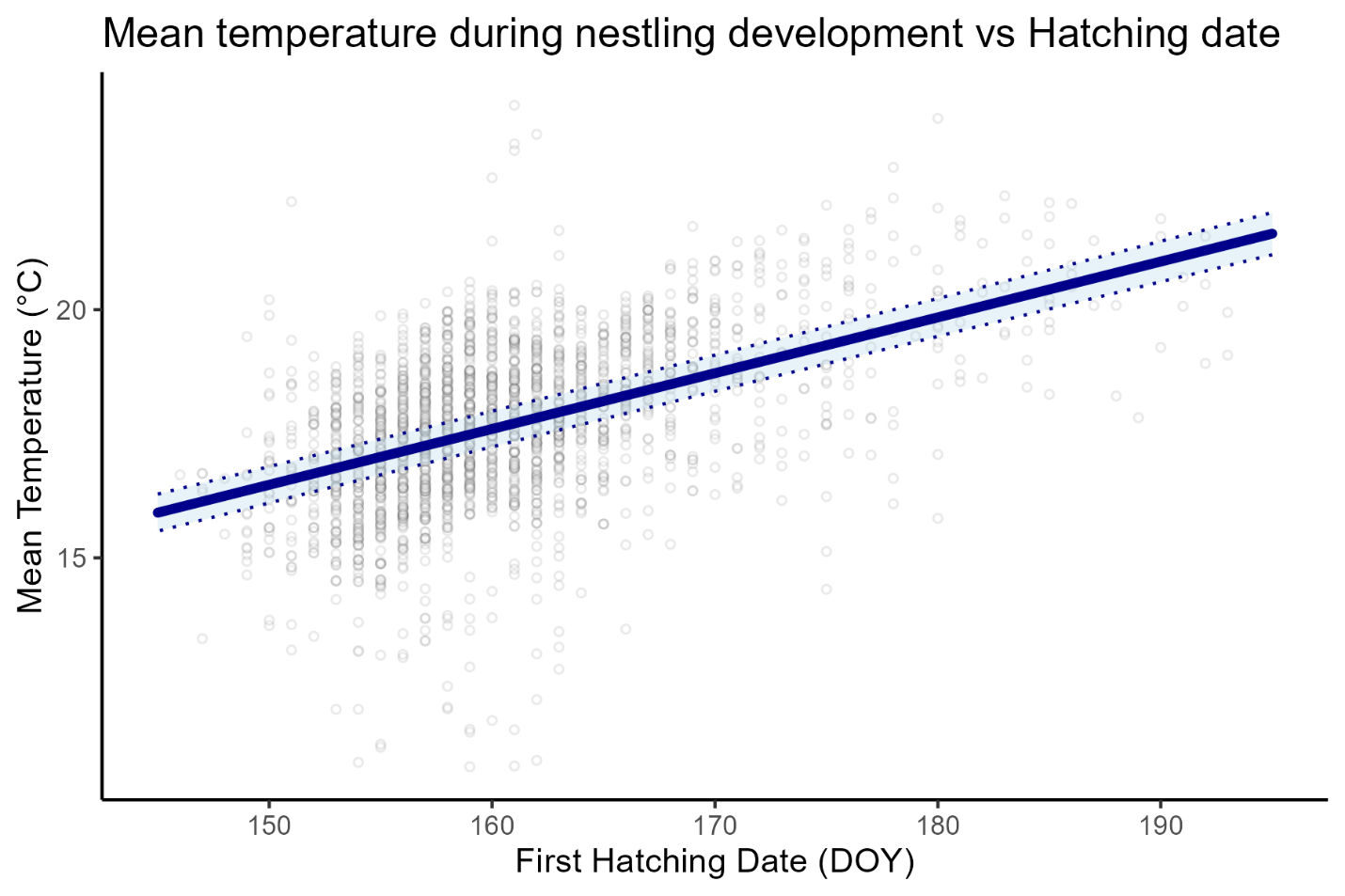

**Figure S2.** *Prediction of mean temperature during nestling development (°C) according to phenology (First hatching date) of a Tree Swallow clutch. 95% Confidence interval in light blue. Raw data in grey dots.*
